# Gentamicin Treatment in a Murine Infection Model Suggests Extracellular *Orientia tsutsugamushi* Contributes to Systemic Dissemination

**DOI:** 10.64898/2026.10.19.760041

**Authors:** Naphat Satapoomin, Chitrasak Kullapanich, Frances Aylward, Manutsanun Inthawong, Kristen Aiemjoy, Carlo Perrone, Tri Wangrangsimakul, Piyanate Sunyakumthorn, Jeanne Salje

## Abstract

*Orientia tsutsugamushi,* an obligate intracellular bacterial pathogen, infects multiple cell types and disseminates via the blood and lymphatic system to affect various organs in human and other mammalian hosts. This bacterium replicates exclusively within living cells; however, it also exists transiently in a metabolically inactive extracellular form. While infected leukocytes are known contributors to the systemic spread of the bacterium, the role of the inactive extracellular form in dissemination has been unclear. In this study, we demonstrate that extracellular bacteria are present in the cell-free plasma fraction of infected human and murine blood. Furthermore, treatment with gentamicin, a membrane-impermeable antibiotic, significantly reduces systemic bacterial growth in a murine infection model. These findings suggest a potential role for the extracellular form of the bacterium in the spread of infection and raise the possibility of exploring the use of gentamicin to augment conventional antibiotic therapy during scrub typhus infection.

## Introduction

*Orientia tsutsugamushi* (Ot) is an obligate intracellular bacterium in the order Rickettsiales, family Rickettsiaceae, that causes the mite-borne human disease scrub typhus^1–3^. This life-threatening disease has a median mortality rate of 6% in untreated cases^4,5^ and is a leading cause of febrile illness in many rural parts of Asia^5–9^. Related species have recently been described in the Middle East^10^ and South America^11,12^, suggesting a more global distribution than previously appreciated.

The overall intracellular infection cycle of Ot has been studied in a range of mammalian cell types using *in vitro* infection models. Ot enters cells through induced clathrin-mediated endocytosis^13^ or macropinocytosis^14^ but exits the endolysosomal pathway shortly after entry^13^ whereupon it replicates directly in the cytoplasm of host cells. This localisation provides direct access to a nutrient rich environment, which is required by this highly reduced bacterium that lacks many genes for metabolite biosynthesis. Ot traffics to the perinuclear region using dynein-dependent motility^15,16^ and undergoes replication in a bacterial microcolony. Following replication and beginning 3-4 days post infection, the bacteria traffic to the cell periphery and bud off the surface of infected cells in a membrane encased extracellular form^17,18^. These extracellular bacteria (EB) are inactive in terms of replication and protein synthesis but can infect naïve host cells^14^. This mode of cell-to-cell transmission differs from that of the closely related sister genus *Rickettsia* species. Whilst cell-free *Rickettsia* are infectious, bacteria from one infected cell can also spread laterally through a tissue using host cell actin to protrude into and penetrate adjacent cells^19^. By contrast, Ot is not known to be able spread directly into adjacent cells without undergoing an extracellular stage, and this is a fundamental difference between the modes of cell-to-cell transfer between these two genera.

Scrub typhus disease initiates with bacterial inoculation at the mite bite site but progresses with dissemination of Ot to multiple organs including the lung, liver, spleen, kidneys and brain^20–22^, where severe infection can result in multiorgan failure and high mortality in untreated disease. During human infection, Ot has been shown to infect vascular endothelial cells in autopsy tissues^23^, as well as dendritic cells and macrophages at the inoculation site in skin biopsies^24^. Murine infection models have further demonstrated rapid dissemination through both the blood and lymphatic systems^21,25,26^, with infected dendritic cells and macrophages contributing to initial bacterial spread to the draining lymph nodes^26^, and protection from humoral immune responses. Recent studies have also highlighted the importance of endothelial activation and vascular inflammation in disease pathogenesis^27^. Consistent with the intracellular nature of circulating Ot, molecular diagnosis of scrub typhus is typically performed using buffy coat samples enriched for white blood cells or whole blood^28–30^, although detection of Ot DNA in serum and plasma samples has also been reported^31^.

Whilst dissemination via infected leukocytes is thought to contribute to the systemic spread of Ot, the potential role of the EB form during infection remained unclear. In particular, it was not known whether EB bacteria were present at detectable levels in plasma, nor whether they contributed to bacterial dissemination in vivo. Here, we investigated the presence and role of the EB form of Ot during animal infection using the membrane-impermeable antibiotic gentamicin, and assessed whether targeting extracellular bacteria could augment conventional antibiotic therapy during scrub typhus infection.

## Results

### Ot is present in the cell free fraction of blood in infected mice and humans

Ot has been shown to bud off the surface of infected cells *in vitro,* but it was not known whether EB form Ot was present in blood during *in vivo* infection. We collected blood from mice that had been infected with 1x10^7^ Ot for 9 days, and human scrub typhus patients, and separated it into a buffy coat fraction containing white blood cells, and a cell free plasma sample. We measured the levels of Ot in both samples using qPCR against the conserved Ot gene *tsa47* (Fig. 1A, B). High bacterial levels were observed in buffy coat samples, as expected, with values ranging from 202-1,173 copies per µl in mice and 9.8-8,686 copies per µl in humans. However, we could also measure low but detectable levels of Ot DNA in plasma samples from infected mice and humans. In total 7/8 mouse plasma samples and 29/64 human patient plasma samples contained detectable Ot DNA with concentrations ranging from 1.8-8.5 copies per µl (mouse) and 2.5-122 copies per µl (human). To confirm that this DNA was associated with bacteria and not with residual mammalian cells, nor free DNA in the plasma samples, we subjected an additional set of 11 human plasma samples to a series of differential centrifugation steps. Where present, bacterial DNA was detectable in fractions corresponding to the bacterial pellet and not mammalian cells (Supp. Fig. 1). One sample gave detectable DNA in both the bacterial pellet and, at a 10x lower level, in the supernatant corresponding to free DNA. Together these data demonstrate that Ot bacteria can be detected in human and murine cell free plasma samples.

**Figure 1.**
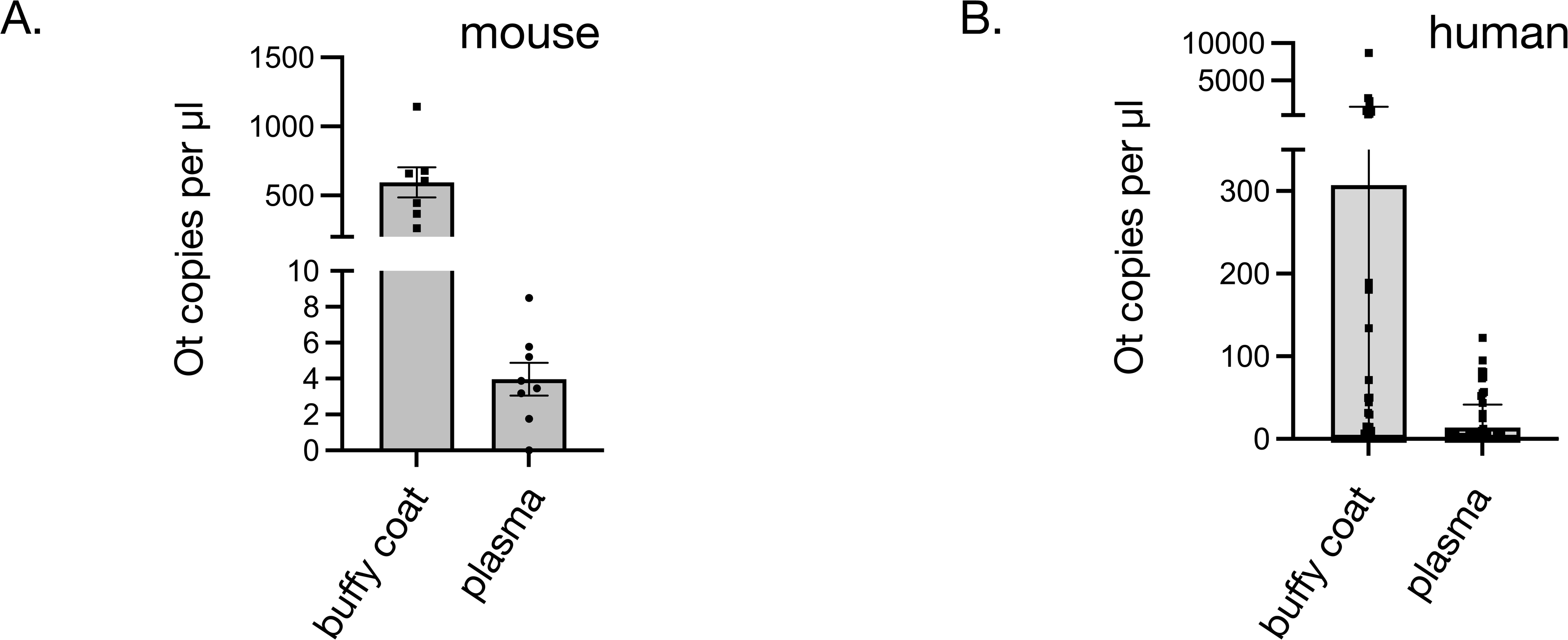
Ot is present in the cell-free fraction of blood in infected mice and humans. A. qPCR analysis showing detectable bacteria in the plasma fraction of mice. Each datapoint corresponds to a single plasma or buffy coat fraction from one mouse (total n = 8). B. qPCR analysis showing detectable bacteria in the plasma fraction of 32 human patients (EXIST study). Each datapoint corresponds to plasma or buffy coat fraction from one patient.

### Gentamicin penetrates extracellular Ot and inhibits its subsequent growth in cultured cells

Gentamicin is a bactericidal aminoglycoside antibiotic that exhibits very low penetration into eukaryotic cells^32–34^ and is not effective as a treatment against murine scrub typhus^35^. These features make it a suitable reagent to determine whether specific inhibition of EB form Ot would affect *in vivo* dissemination. Its effect on different Ot populations *in vitro* had not been reported so we carried out experiments to determine its activity (Fig. 2A). First, we prepared intracellular form (IB) and EB form Ot and exposed them to a range of concentrations of gentamicin for 6 hours. Gentamicin treatment of IB Ot was carried out by treating infected cells, whilst treatment of EB form was carried out on bacteria that had been isolated from the supernatant of infected cells. Following antibiotic treatment, IB bacteria were isolated from host cells by mechanical lysis and centrifugation and both IB and EB Ot used to infect naïve host cells. The infections subsequently proceeded for 1 and 5 days, whereupon the bacterial concentration was determined by qPCR (Fig. 2B-E). A comparison of the bacterial copy number at day 1 and day 5 shows similar growth of IB and EB form Ot in the absence of gentamicin treatment, as shown previously (compare the increase in bacterial number with 0 µg/ml gentamicin between Fig. 2B and C, and Fig 2D and E). Gentamicin treatment had no effect on the ability of IB form Ot to replicate *in vitro* (Fig. 2B, C). By contrast, EB Ot was inhibited by gentamicin treatment in a dose-dependent manner (Fig. 2D, E). This data shows that gentamicin inhibits the growth of EB form Ot but not IB form Ot.

**Figure 2.**
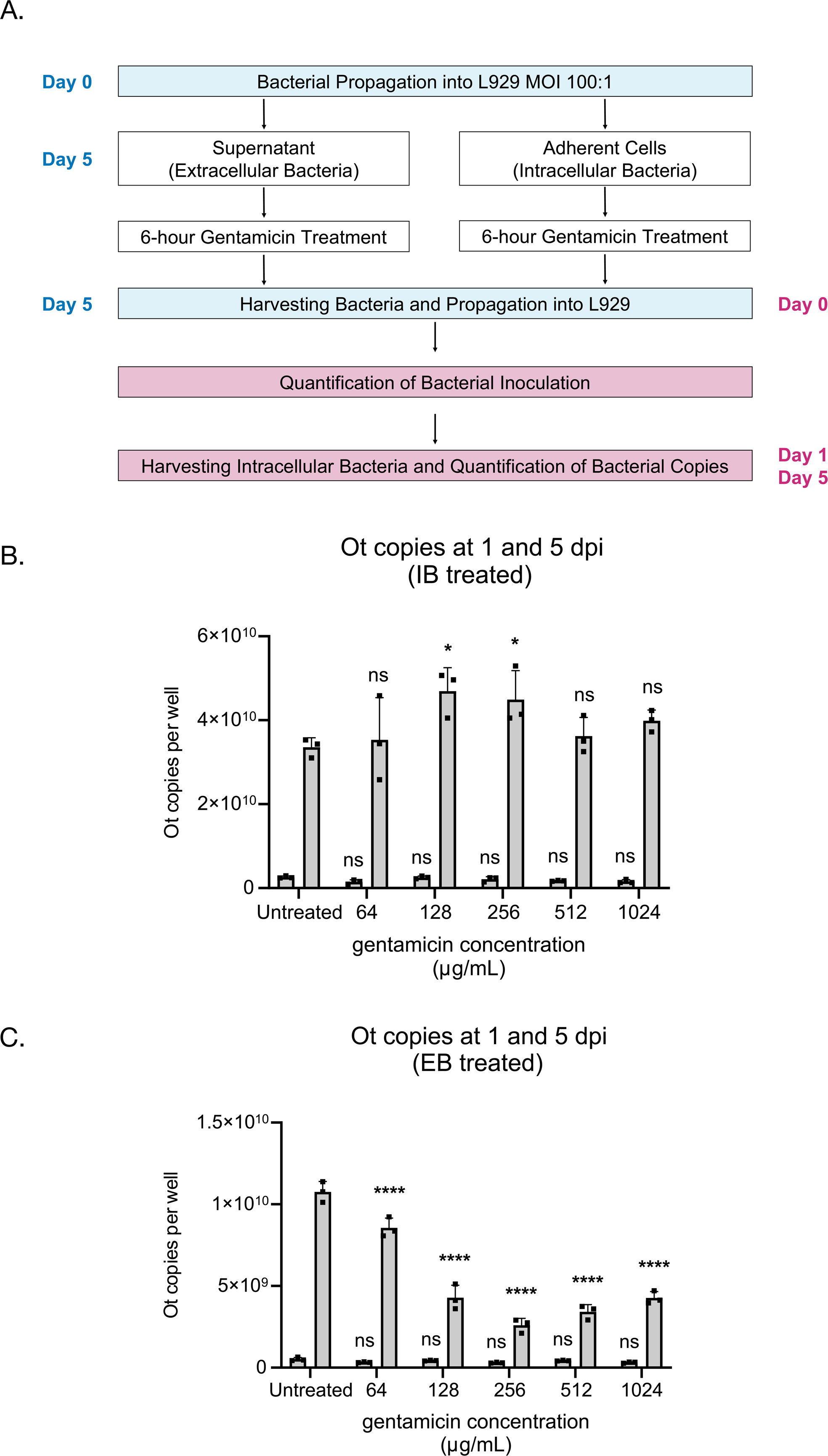
Gentamicin penetrates into extracellular Ot and inhibits its subsequent growth in cultured cells. A. Overview of experiment testing the effect of gentamicin on intracellular (IB) and extracellular (EB) bacteria *in vitro.* An expanded version of this protocol is given in Supp. Fig. 2. Ot was exposed to gentamicin either whilst inside L929 cells (IB treated: B) or after being isolated from host cells (EB treated: D). The treated bacteria were then infected onto L929 cells and grown for 1 day or 5 days and the bacterial number was quantified by qPCR. Grouped bar graphs show bacterial number after 1 day on the left and after 5 days on the right. Gentamicin treatment inhibited the growth of bacteria exposed to gentamicin when they were outside host cells (C), but not those that had been inside host cells (B) at the time of drug treatment. One-way ANOVA with Dunnett’s multiple comparisons was used to calculate the statistical significance of the change in growth upon treatment compared to untreated control. **** < 0.0001.

To further demonstrate that gentamicin could penetrate EB Ot, we prepared EB Ot and exposed it to a fluorescently-conjugated gentamicin derivative, Gentamicin-Texas Red. Microscopy of bacteria exposed to this conjugate for 24 hours demonstrated uptake of gentamicin into 54.8 % (s.d. 8.05%) of EB Ot (Fig. 3), demonstrating that this antibiotic can enter the bacteria in its extracellular form.

**Figure 3.**
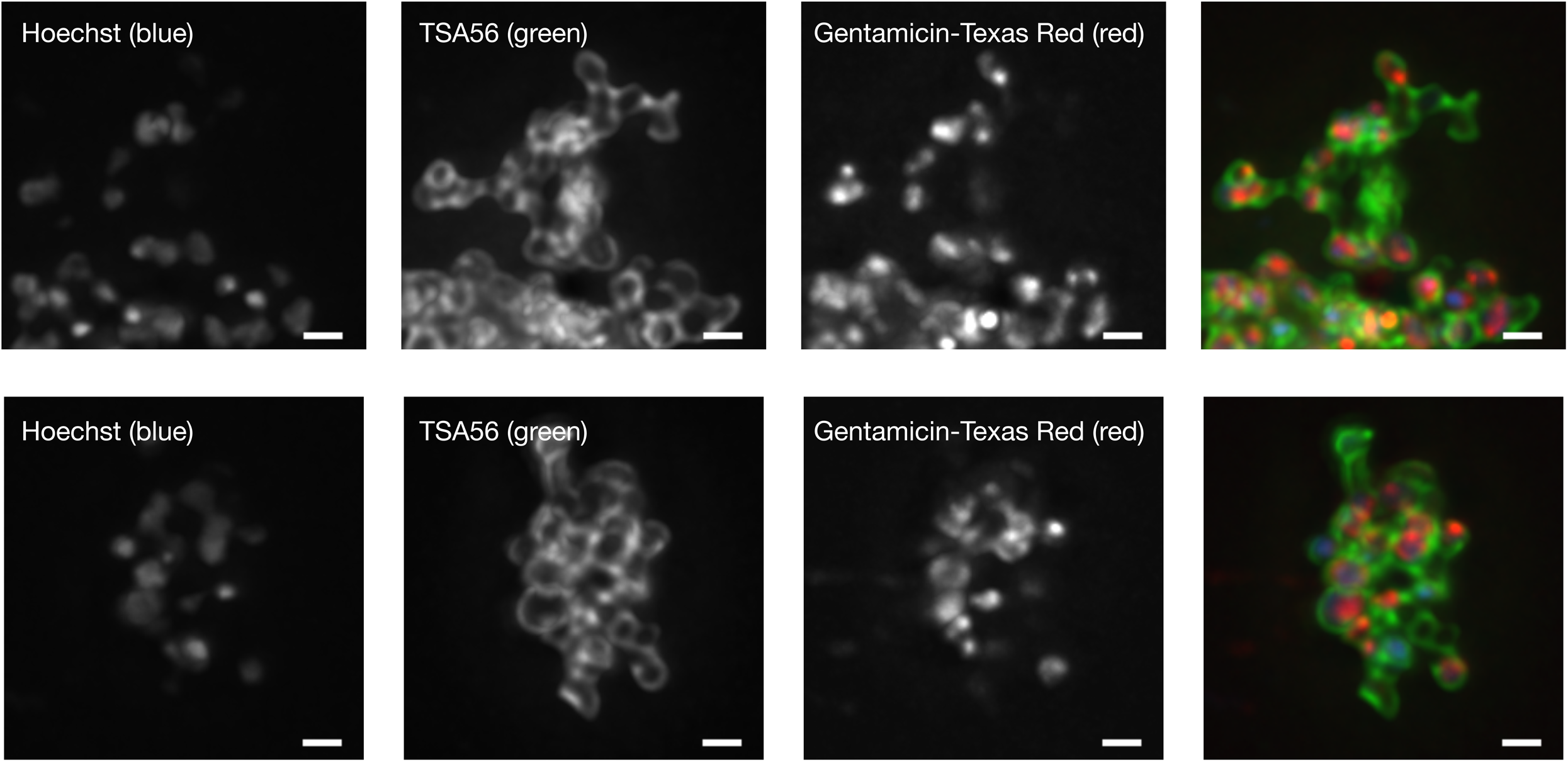
A fluorescent derivative of gentamicin penetrates extracellular Ot *in vitro*. Fluorescence micrograph of extracellular bacteria incubated with fluorescent gentamicin showing penetration of this drug conjugate into bacteria. Bacteria are labelled using an antibody against the surface protein TSA56. Green = TSA56 antibody, Red = Gentamicin-Texas Red. Scale bar = 10 µm.

### Gentamicin treatment reduces bacterial dissemination to various organs in a murine infection model

We sought to determine whether EB form Ot plays a role in bacterial dissemination *in vivo* by selectively reducing this population using gentamicin treatment and measuring bacterial levels during infection (Fig. 4A). We inoculated C57Bl/6NJcl mice with 1x10^7^ genome copies, corresponding to 3.3x10^6^ focus forming units, of Karp strain Ot using intradermal inoculation. Intradermal inoculation was used to ensure that bacteria detected in blood samples originated from dissemination from infected tissues rather than from the initial inoculum, and a sublethal infection model was used to allow consistent quantification of bacterial burden without confounding effects from severe disease or variable mortality. Infections proceeded without antibiotic treatment for 5 days to allow bacteria to reach different target organs. At 6 days post infection, gentamicin was administered by intravenous inoculation at low dose (6 mg per kg) or high dose (24 mg per kg). Control groups lacking gentamicin, with only gentamicin (no Ot), or with only buffer, were included in this experiment. Infections proceeded until 9 days post infection whereupon the bacterial load in blood and tissues was determined by qPCR (Fig. 4B-C). Analysis of different blood samples showed a small but not statistically significant reduction in the bacterial level in both buffy coat and plasma fractions upon gentamicin treatment (Fig. 4B). By contrast, significant reductions in bacterial copy number were observed in brain, kidney, lung and spleen, with a small and not significant reduction observed in the liver (Fig. 4C). Overall, the results were similar between low and high dose of gentamicin, although slightly fewer bacteria were detected in the brain, kidney and spleen in the presence of lower gentamicin compared with higher. Together these data show that Ot blood levels are not significantly reduced by gentamicin treatment in a murine infection model but the levels in brain, kidney, lung and spleen are significantly reduced.

**Figure 4.**
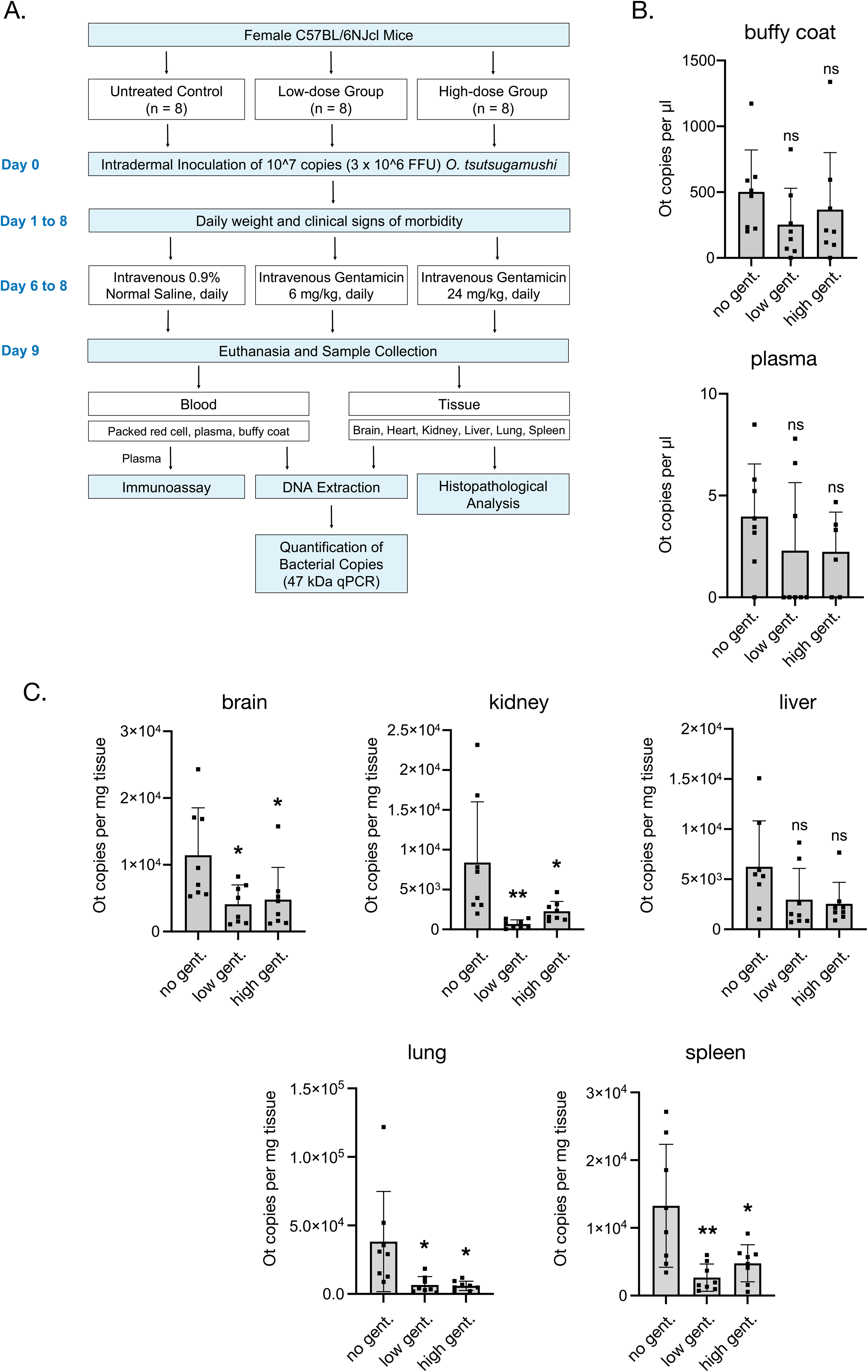
Gentamicin treatment reduces bacterial dissemination to various organs in a murine infection model. A. Schematic showing the experimental design for an *in vivo* non-lethal murine infection model where C57Bl/6NJcl mice were intradermally inoculated with 1x10^7^ copies (3.3x10^6^ focus forming units) of Ot strain Karp. Beginning at 6 days post infection gentamicin was administered intravenously at a low dose of 6 mg per kg, or high dose of 24 mg per kg, once daily. Eight animals were used in each experimental group and the entire experiment was carried out two independent times. Similar results were obtained in both experiments and the results of one repeat are shown here. B. Quantification of Ot bacterial copy number in buffy coat and plasma blood fractions at 9 days post infection. Mean and SD is shown. No statistically significant differences were measured based on one-way ANOVA analysis. C. Quantification of Ot bacterial copy number in brain, kidney, liver, lung and spleen tissues at 9 days post infection. Mean and SD is shown. Statistical significance was calculated using one-way ANOVA analysis followed by Dunnett’s multiple comparisons test. * p ≤ 0.05, ** p ≤ 0.01, ns p > 0.05.

### Gentamicin-induced reduction in bacterial tissue load does not lead to detectable differences in disease

We asked whether the observed reduction in bacterial load in brain, kidney, lung and spleen resulting from gentamicin treatment would lead to improved disease outcomes in this *in vivo* infection model. We measured weight change during the infection presented in Fig. 4 and observed no significant differences between any groups (Fig. 5A). Note that the experiment in this study was deliberately carried out using a low bacterial dose and intradermal inoculation, leading to mild and non-lethal disease, as this would better reveal differences in bacterial load upon gentamicin treatment. Consistent with this, there was no weight loss observed in the Ot group in the absence of gentamicin and therefore no differences were expected between groups. We quantified clinical score using loss of appetite, lethargy and ruffled fur and markers of disease. Mild disease was observed on day 9 for the Ot infected groups, and this was not affected by gentamicin treatment (Fig. 5B). Finally, we asked whether the reduced bacterial load in tissues resulting from gentamicin treatment would affect the inflammatory response to infection. We measured the cytokine profile of plasma samples taken 9 days post infection. Ot infection led to an induction of IFN-γ, IL-6, MCP-1, as has been described previously^36,27,37–41,22^, but this induction was not affected by gentamicin treatment (Fig. 5C). High dose gentamicin treatment in the absence of Ot led to a small increase in IL-6 levels, likely as a result of liver toxicity^42^. Overall, these data indicate that gentamicin treatment does not affect disease outcome under these conditions in this murine infection model.

**Figure 5.**
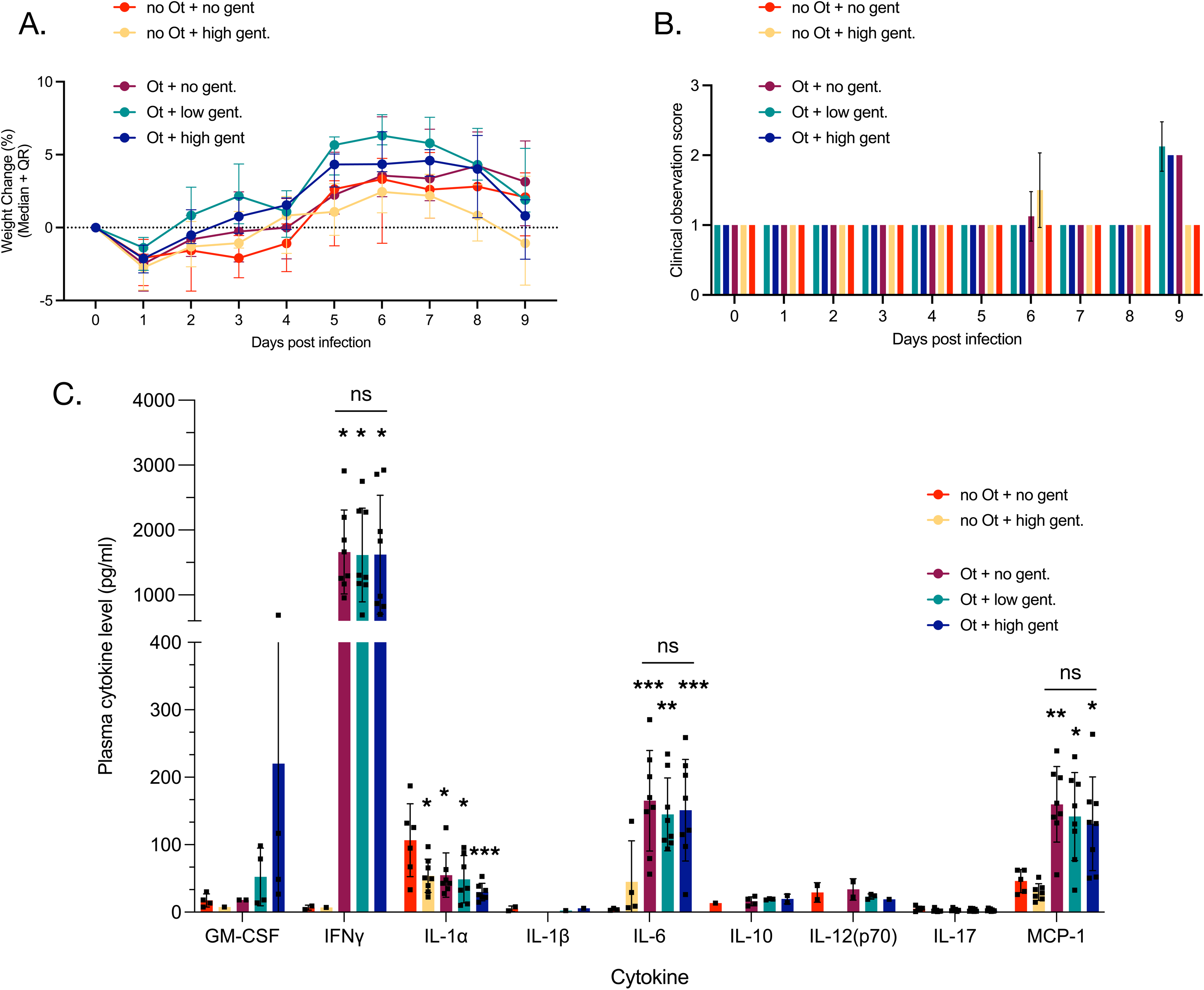
*in vivo g*entamicin treatment does not significantly change disease outcomes in a murine infection model. Clinical outcomes and inflammatory response in the same murine infection experiment described in Fig. 4. A. shows the weight change or each group at each time point compared to day 0. Median and quartile range is shown. Statistical analysis was carried out using two-way ANOVA and no statistical significances were determined. B. Clinical observations were made daily, and scored based on loss of appetite, lethargy and ruffled fur appearance. The mean and SD for each group are shown. No statistically significant differences were measured based on two-way ANOVA analysis. C. Relative levels of cytokines in murine plasma samples at 6 days post infection as measured by cytokine analysis. Mean and SD is shown, together with results of one-way ANOVA analysis with Dunnetts multiple comparisons where each group is compared with the no Ot + no gent. control group (asterix above bars). The same statistical test was used to look for differences between Ot +/- gentamicin groups and no significant differences were detected for any cytokine (ns across bars). * p ≤ 0.05, ** p ≤ 0.01, *** p ≤ 0.001, ns p > 0.05.

## Discussion

Whilst dissemination via infected leukocytes plays a major role in Ot spread^21,25,26^, the potential role of cell free EB bacteria was unknown. It was not known if it was present in detectable levels in plasma, nor whether it was important for dissemination. Here, we set out to assess whether EB was detectable in *in vivo* infection, and to assess the role in systemic infection.

We show that the antibiotic gentamicin selectively reduces viability of EB form Ot *in vitro*, and that gentamicin treatment of infected mice leads to a reduction of Ot in lungs, brains, kidneys and spleens of mice. There are two possible factors that might drive this reduction. First, the gentamicin treatment may affect the viability of extracellular Ot in plasma, limiting the bacteria’s ability to disseminate systemically around the body. Note that these bacteria may be EB form Ot that has budded off the surface of intact infected cells, or they may derive from IB Ot released by damaged infected cells. Second, gentamicin may affect the local dissemination of Ot between cells within an organ, through its presence in interstitial fluid. This interpretation is consistent with our observation that gentamicin treatment has no significant effect in the liver, an organ with a compact architecture and little interstitial fluid. Our data shows that extracellular Ot plays a role in dissemination but cannot currently distinguish the relative importance of Ot EB in local and systemic dissemination.

The finding that Ot EB plays a role in dissemination has potential implications for vaccine development, because Ot EB can potentially be accessed by antibodies and complement in a way that is not possible when bacteria are encased within host cells. This is consistent with the fact that Ot stimulates a strong antibody response and the fact that passive immunization experiments have been successful in scrub typhus research^43,44^, although cellular immunity – that would not target EB Ot – is known to be essential for immunity in scrub typhus infection^45–47^. We observe that only ∼50% of bacteria take up fluorescent gentamicin (Fig. 3), and that EB gentamicin treatment does not lead to complete loss of growth of these bacteria (Fig 2D, E). This suggests that not all EB bacteria in the population can be targeted by gentamicin. This may result from different physiological states of different individual EBs, or from the lingering presence of encasing host plasma membrane formed during bacterial budding. Future studies into these processes may lead to improved methods to target the RB population during infection using a combination of antibodies and antibiotics.

We found that gentamicin treatment reduced bacterial load in tissues but did not affect disease outcomes under these experimental conditions. One potential explanation is that this is due to the nonlethal murine infection model that was selected for this study, as well as the relatively short time during which gentamicin was administered. However, it is surprising that the magnitude of the inflammatory response is consistent between gentamicin treatment and control groups, as cytokine levels were previously shown to differ significantly between strains of different virulence. Our data suggests that the magnitude of the inflammatory response in plasma is not directly correlated to the bacterial load in lungs, brain, kidney and spleen.

We sought to design an experiment that would selectively target Ot EB, and we used the non-permeable antibiotic gentamicin to achieve this. One limitation of our study is the fact that gentamicin has been shown to have low levels of activity against some other intracellular bacteria suggesting low levels of penetration into eukaryotic cells^48–52^, although this is thought to primarily occur through uptake into the endosomal pathway^49^ and thus be less effective against cytoplasmic bacteria such as Ot. Our *in vitro* measurements show no activity against intracellular Ot, but it remains possible that residual activity against Ot IB was achieved *in vivo* and this is a limitation of our study. The observation that gentamicin induced only a small and non-significant reduction in bacterial load in the liver, where there is low interstitial fluid, compared with other organs, supports an interpretation that any intracellular effect is minimal. An additional limitation of our study is the fact that the use of a non-lethal murine model limits the ability to assess the impact of gentamicin treatment on disease.

Our data show that gentamicin treatment in a sublethal intradermal murine infection model reduces bacterial burden in multiple tissues but does not prevent the development of clinical disease. This contrasts with other antimicrobials that have demonstrated protective efficacy in vivo, including the RNA polymerase inhibitor CorA^53^, chloramphenicol, and ciprofloxacin^35^. As gentamicin does not target the IB form of Ot, it would not be expected to fully clear infection or prevent disease progression. However, alternative treatment regimens or the use of intravenous or intraperitoneal infection models may enhance bacterial clearance and warrant further investigation. It is also possible that combining gentamicin with antimicrobials active against IB Ot, such as doxycycline, could improve in vivo bacterial clearance, analogous to the reported superiority of combined azithromycin/doxycycline therapy over monotherapy in severe scrub typhus cases^54^.

In summary, we have shown that Ot is present in the cell free plasma fraction of infected humans and mice, and that preferential depletion of the cell free EB form of Ot through gentamicin treatment leads to a reduction in bacterial load in certain tissues in mice. This advances our understanding of bacterial dissemination during disease and has implications for the design of scrub typhus vaccines and improved antibiotic treatment regimens.

## Materials and Methods

### Bacterial Propagation

*Orientia tsutsugamushi* strain Karp was propagated in mouse fibroblast cell line NCTC clone 929 (L929) purchased from ATCC (Cat. CCL1). L929 cells were seeded in T75 flask containing DMEM (Gibco 11965092) supplemented with 10% FBS (Sigma F7524) at a density of 1.2 × 10^6^ cells per flask and incubated at 37°C with 5% CO_2_ until confluence was reached. A frozen bacterial stock was thawed and added directly to the L929 at a multiplicity of infection (MOI) of 100. After 3 hours incubation, bacteria were removed, and fresh DMEM medium supplemented with 10% FBS was added to each flask. On day 5 post-infection, the medium was removed followed by washing with Phosphate Buffered Saline (PBS) once and the cells were then scraped from the flask. Cells were resuspended in DMEM medium and then transferred to a 2 ml Safe-Lock tube and homogenised using a bead-mill homogeniser at the speed of 8 for 1 minute. The suspension was then centrifuged at 300 xg for 5 minutes to separate the host cells. Supernatant was collected then centrifuged at 20,000 xg for 3 minutes to pellet Ot bacteria. The bacterial pellet was resuspended with Sucrose-Phosphate-Glutamate (SPG) buffer and stored at -80 °C ready for use in experiments.

### Mouse Experiment

All animal research was approved by the AFRIMS Animal Care and Use committee (Protocol number 24-03), and conducted in compliance with Thai laws, the U.S Animal Welfare Act, and guidelines from the U.S. Department of Agriculture (USAD), Office of Laboratory Animal Welfare (OLAW) and U.S. Department of Defence guidelines. The study adhered to the *Guide for the Care and Use of Laboratory Animals* (8^th^ Edition, NRC publication) and was conducted at the AAALAC-accredited AFRIMS facility in Bangkok, Thailand.

Female C57BL/6NJcl mice (6-9 weeks old) were sourced from Nomura Siam International Co., Ltd (Bangkok, Thailand) and housed under specific pathogen-free (SPF) conditions in an animal biosafety level 2 (ABSL2) facility. Mice were co-housed (4 mice/cage, 2 cages/group) in ventilation polycarbonate microisolator cages with standard feeding and watering system maintained at 21 °C ± 1, with relative humidity between 30–70%.

On Day 0, mice (n= 8 per group) were chemically immobilized with intramuscular ketamine (5-20 mg/kg) and intradermally inoculated at a shaved dorsal site with 50 µl of *O. tsutsugamushi,* Karp strain (10^7^ genomic DNA; equivalent to 3.3 x 10^6^ FFU) or mock control using a 30G ½” syringe (Ultra-Fine II, Becton Dickinson). On days 6 to 8 post-inoculation, mice received once-daily intravenous gentamicin (VESCO Pharmaceutical Co., Ltd) at either low (6 mg/kg) or high (24 mg/kg) dose or normal saline 0.9% (General Hospital Products Public Co., Ltd) as a control. Mice were weighed daily, and clinical signs of morbidity were monitored and scored throughout the 9-day experimental period. Animals exhibiting signs of severe morbidity or >15% weight loss from baseline were humanely euthanized. At the study end point (Day 9), blood and tissues (brain, heart, lungs, kidneys, spleens, and liver) were collected for bacterial quantification and histopathological analysis. Mice were humanely euthanized using CO_2_ inhalation with gas flow of 2 l/min at 15 psi CO_2,_ maintained for at least 5 min after the animal ceased breathing. Death was confirmed by physical examination (the absence of a heartbeat) and ensured with an adjunctive physical method such as cervical dislocation or exsanguination.

### Focus Forming Unit (FFU) Measurement of Bacterial Samples used in Mouse Studies

One aliquot of Ot Karp strain prepared for inoculation in the mouse model was propagated into L929 cells seeded in 8-well chamber slide at a starting MOI of 10^4^. Propagation at 10^-1^, 10^-2^, 10^-3^ and 10^-4^ dilutions was also performed. After 6 hours of infection, cells were fixed with 4% formaldehyde for 10 minutes at 4°C. Cells were then permeabilised using absolute ethanol for 1 hour on ice. After 1 hour, cells were washed with PBS and further permeabilised using 0.5% Triton X for 30 minutes on ice. Cells were then washed with PBS-T (PBS with 0.05% Tween 20) and incubated with primary antibody (rat monoclonal antibody against TSA56 1:200 dilution) at 4°C overnight. After primary antibody was removed, cells were then incubated with secondary antibody (goat anti-rat Alexa Fluor 488 ThermoFisher A11006 at 1:1000 dilution), supplemented with nuclear stain Hoechst (1:2000 dilution) at 37°C for 1 hour. Secondary antibody was removed samples were washed with PBS-T then mounted using Prolong^TM^ Gold Antifade Mountant (ThermoFisher P36930). Samples were then visualized using a Zeiss Observer Z1 LSM700 confocal microscope with an HBO 100 illuminating system equipped with a Plan-APOCHROMAT 63x/1.4 oil objective lens (Carl Zeiss, Germany). Phase contrast (T-PMT, DIC), laser line 405 nm, and laser line 488 nm were used for image acquisition using ZEN 2011 SP7 FP3 (version 14.0.0.0) software. The number of bacteria was counted from 3 frames per dilution and averaged as a final FFU value.

### Quantification of Bacteria in Human Clinical Samples

A total number of 32 human plasma and 32 buffy coat samples were obtained as a part of the Eschar Investigations in Scrub Typhus (EXIST) study (Clinicaltrials.gov identifier NCT02915861). Whole blood samples were originally collected from patients admitted at Chiangrai Prachanukroh Hospital, Chiangrai province, Thailand. Patients presented febrile illness, eschar, and tested seropositive for scrub typhus IgM RDT (InBios International, Inc. USA). The volume of 200 μl of each human plasma and buffy coat sample were subjected to DNA extraction using the DNeasy Blood and Tissue Kit for DNA Isolation (QIAGEN 69504) following manufacturer’s instruction and bacterial quantification using qPCR targeting 47 kDa gene of *O. tsutsugamushi* as described in previously deposited protocol^55^. The primer pair and TaqMan probe used were as follows: 47 kDa FW (5’-TCCAGAATTAAATGAGAATTTAGGAC-3’), 47 kDa RV (5’-TTAGTAATTACATCTCCAGGAGCAA-3’), and 47 kDa probe (5’-FAM-TTCCACATTGTGCTGCAGATCCTTC-TAMRA-3’). The qPCR mixture consisted of 1 μL extracted DNA template, 1X PCR Probe Mix Lo-ROX (PCR Biosystems PB20.21), 0.1 μM of each forward and reverse primer, 0.2 μM TaqMan probe, and nuclease-free water to make up a total volume of 20 μL. CFX96 Real-time PCR Detection System and CFX Manager Software (Bio-Rad) were used to perform real-time PCR using the following the conditions: initial denaturation at 95°C for 2 minutes, 45 cycles of 95°C for 15 seconds and 60°C for 30 seconds. DNA copy numbers were calculated by comparison with a standard curve constructed from a serial dilution of 47 kDa standard in the range of 10^8^ to 10^1^ copies.

Samples were collected by trained research nurses with valid good clinical practice (GCP) certification. The study was approved by the Mahidol Faculty of Tropical Medicine EC (ref. no: T-MEC 16/065), the Chiangrai Prachanukroh Ethics committee (CRH-EC), and the Oxford Tropical Medicine Ethics Committee (OxTREC Ref. no. 46-15). All participants provided informed consent and understanding the pathophysiology of scrub typhus was within the remit of the study and included in the participant information sheet.

Additional bacterial copy quantification was carried out on 11 human plasma samples. These samples were subjected to additional two-step centrifugation step to separate host cells from bacterial cells before DNA isolation. A total volume of 100 μL of plasma was centrifuged at 300 xg for 5 minutes to pellet the host cells. The supernatant was collected and centrifuged at 20,000 xg for 3 minutes to pellet the bacterium. We collected 10 μl of sample from each step: 1) before any centrifugation, 2) pellet from low-speed centrifugation, 3) supernatant after high-speed centrifugation, and 4) pellet from high-speed centrifugation to determine the presence of bacteria in each fraction. All samples were subjected DNA isolation using alkaline lysis method and bacterial quantification was carried out by 47 kDa method described above.

### Quantification of Bacteria in Mouse Samples

Plasma, buffy coat, and mouse tissue (brain, heart, kidney, liver, lung, spleen) samples were subjected to DNA extraction using DNeasy Blood and Tissue Kit for DNA Isolation (QIAGEN 69504) following the recommended protocol by the manufacturer. The volume of 100 μl of each plasma and buffy coat sample was used for DNA isolation. The final bacterial copies were calculated using the original volume of sample used and reported in ‘copies per μl’ format. Similarly, all tissue samples were dissected and weighed to not exceed 20 mg, as suggested by the manufacturer for spin column capacity. The weight of tissue was then used to calculate and report final bacterial concentration in ‘copies per mg’ format. Bacterial copies were quantified using qPCR targeting 47 kDa gene of Ot as described above.

### Immunoassay

Mouse plasma samples collected at euthanasia were assessed using the MILLIPLEX multiplex assay (MCYTOMAG-70K, Merch Millipore) following the manufacturer’s instructions. A total of nine analytes were measured, including GM-CSF, IFN-γ, IL-1α, IL-1β, IL-6, IL-10, IL-12(p70), IL-17, and MCP-1. Quantification of these cytokines was performed on a MAGPIX (Luminex) analyser and the Belysa immunoassay software 1.2.2 was used for data collection.

### *In vitro* Gentamicin Assay

L929 cells at a density of 3 × 10^6^ cells were seeded in six T175 flasks containing DMEM (Gibco 11965092) medium supplemented with 10% FBS (Sigma F7524) and incubated at 37°C with 5% CO_2_. After 2 days, Ot Karp strain was propagated into L929 at an MOI of 100. Bacteria were removed after 3 hours of incubation at 35°C with 5% CO_2_ and fresh DMEM medium with 10% FBS was added to each flask. At day 5 post-infection, the supernatants (extracellular bacteria) were collected in 50 ml centrifuge tube. Gentamicin (Sigma G1914) was prepared on the day by dissolving in water to achieve a working concentration 64 mg/ml, filter-sterilised using 0.22 μM filter. The calculated volume of gentamicin was added to DMEM medium at a concentration of 0, 64, 128, 512, and 1024 μg/ml to both supernatants collected (extracellular bacteria) and adherent cells (intracellular bacteria). The chosen concentration range was benchmarked against an estimated concentration of gentamicin in the mouse infection model. The benchmark concentrations of 232.99 μg/mL and 931.96 μg/mL were calculated from gentamicin doses (low dose and high dose groups) divided by the average total blood volume collected at end point. Infected cell line was incubated with gentamicin for 6 hours at 35°C with 5% CO_2_. Supernatant was transferred to 50 ml tube and centrifuged at 20,000 xg at 4°C for 15 minutes to pellet extracellular bacteria (EB). The adherent cells in each T175 flask were scraped and transferred to 2 mL Safe-Lock tubes and homogenised using Bullet Blender Homogeniser for 1 minute at the speed of 8. The suspension was then centrifuged at 300 xg for 5 minutes at room temperature to separate host cells. Supernatant was collected and centrifuged at 20,000 xg for 3 minutes at room temperature to pellet intracellular bacteria (IB). The EB and IB pellets were resuspended with 2 mL of DMEM medium supplemented with 10% FBS. Subsequent infection of Ot Karp with and without gentamicin treatment was performed by adding 300 μl of EB and IB suspension to 24-well plates containing L929 and incubated at 35°C with 5% CO_2_. Bacterial cells were harvested at day 1 and 5 post-infection. The DMEM medium in the 24-well plates was removed. Alkaline lysis buffer was to each well at the volume of 200 μl and the plates were covered with adhesive foil and boiled at 95°C for 30 minutes to isolate DNA. The cell lysates (3 μl from each well) were then used as a template for 47 kDa qPCR to quantify bacterial copy. The final bacterial copy was reported as copy per μl calculated from the original volume of cell lysates. Statistical analysis was performed using GraphPad Prism 10.4.1.

### *In vitro* assessment of gentamicin effect on extracellular bacteria (EB) using fluorescent-labelled gentamicin

L929 cells were seeded in two T25 flasks at 40% confluence and infected the following day with Ot strain Karp at an MOI of 200. On day 6 post-infection, supernatant was filtered through a 5 µm Millex SV Filter Unit (Merck Millipore, SLSV025LS) to remove host cells and spun at 14,000 xg for 5 minutes to obtain EB. EB were resuspended in DMEM with 5% FBS containing 64 µg/mL Gentamicin Texas Red (GTTR) (AAT Bioquest, AATB24300). This solution was incubated at 36°C with 5% CO_2_ for 18 hours, then spotted onto slides which were previously coated in poly-D-Lysine (Sigma-Aldrich, A-003-E) at 100 µg/mL for 1 hour and washed twice with PBS (Sigma-Aldrich, D8662). Samples were then fixed with 4% PFA (Thermo Fisher Scientific, 28906) for 10 minutes in the dark at room temperature. Slides were washed once with PBS and kept in the dark in PBS at 4°C prior to staining. EB were labelled overnight at 4°C in PBS containing 1 mg/mL Bovine Serum Albumin (Sigma-Aldrich, A9647) and Rat anti-TSA56 antibody (In-house) at a concentration of 1:200. Samples were then washed twice with PBS and incubated at 37°C for 1 hour in PBS containing Goat anti-Rat 488 antibody (Thermo Fisher Scientific, A21434) at a 1:500 concentration and Hoechst 33342 at 10 µg/mL (Thermo Fisher Scientific, H1399). Samples were washed twice with PBS, then mounted using in-house mounting media. Samples were imaged using a Nikon Ti2-E inverted microscope with a pE4000 Universal Fluorescence illumination system equipped with a 100XH Oil/1.45 CFI Plan-APOCHROMAT x λ objective lens (Nikon Europe B.V) and LED-DAPI-A-2360A, LED-FITC-A-2745A, and LED-TRITCA filter cubes (32mm). 3D deconvolution of Z-stacks was performed on Nikon NIS-Elements AR Version 5.42.04, followed by Maximum Intensity Projection. Total Ot and GTTR-positive Ot were counted manually from 4 images containing >180 EB Ot per image.

## Acknowledgements

This work was supported by a Wellcome Trust Senior Research Fellowship (224277/Z/21/Z) which contributed the salary of JS, CK and NS. FA was supported by a BBSRC DTP PhD studentship. The funders had no role in study design, data collection and analysis, decision to publish, or preparation of the manuscript. Disclaimer: Material has been reviewed by the Walter Reed Army Institute of Research. There is no objection to its publication. The opinions or assertions contained herein are the private views of the author, and are not to be construed as official, or as reflecting true views of the Department of the Army or the Department of Defense. Research was conducted under an IACUC-approved animal use protocol in an AAALAC International - accredited facility with a Public Health Services Animal Welfare Assurance and in compliance with the Animal Welfare Act and other federal statutes and regulations relating to laboratory animals.

We are grateful for Nicholas Day, MORU, for insightful discussions on this work.

