## Supplementary Material for "Gentamicin Treatment in a Murine Infection Model Suggests Extracellular *Orientia tsutsugamushi* Contributes to Systemic Dissemination"

A.

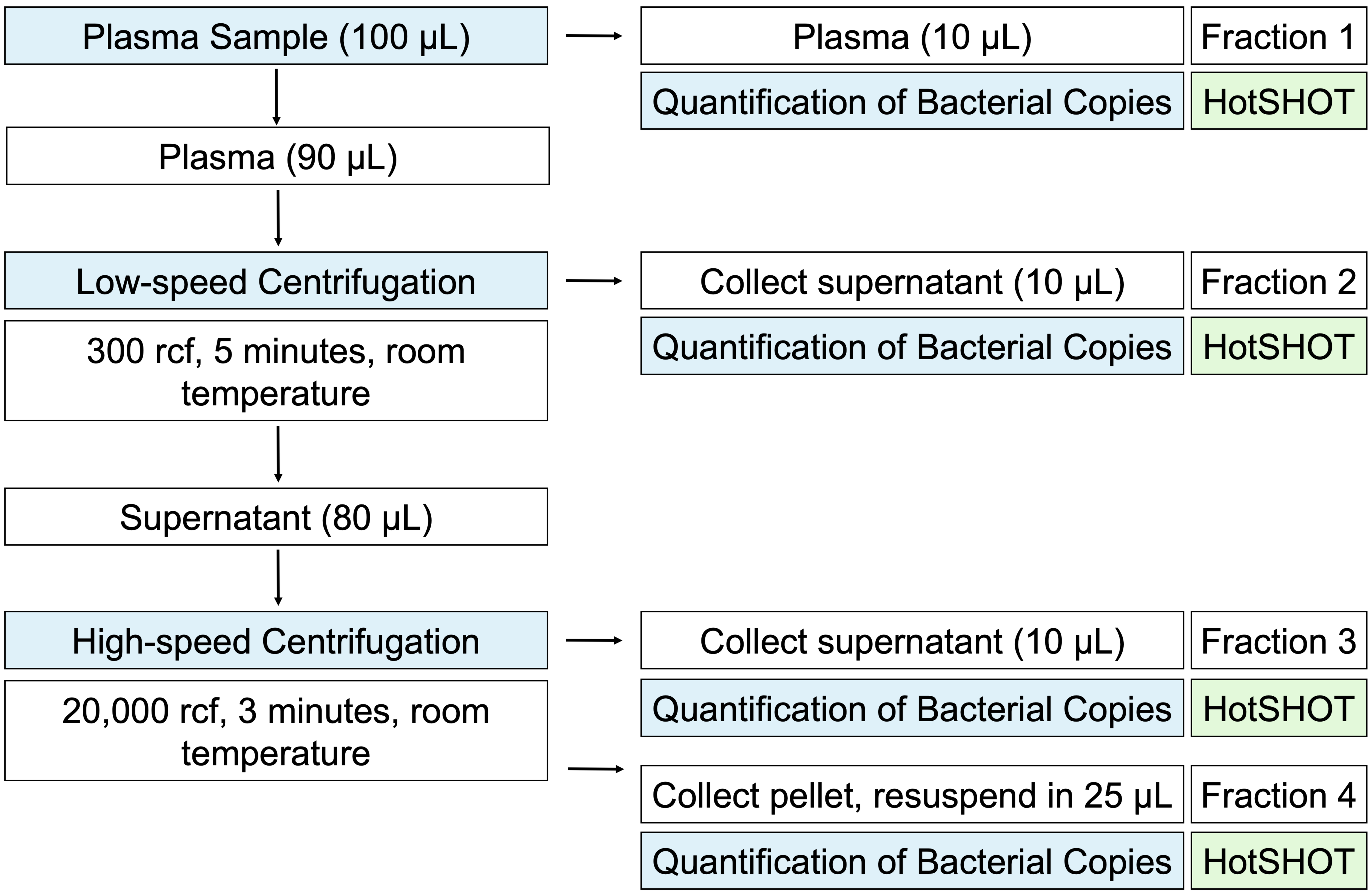


B.

| Sample | Disease Duration (days) | Ot copies/100µL (47 kDa) | | | |
|---|---|---|---|---|---|
| | | Fraction 1 | Fraction 2 | Fraction 3 | Fraction 4 |
| QES-60-002-W0-PE | 3 | Not detectable | Not detectable | Not detectable | 1.12E+00 |
| QES-60-007-W0-PE | 3 | 3.41E+00 | Not detectable | 3.55E+00 | 2.20E+01 |
| QES-60-027-W0-PE | 8 | Not detectable | Not detectable | Not detectable | Not detectable |
| QES-60-040-W0-PE | 10 | Not detectable | Not detectable | Not detectable | 1.04E+00 |
| QES-61-016-W0-PE | 5 | Not detectable | Not detectable | Not detectable | Not detectable |
| QES-61-018-W0-PE | 4 | Not detectable | Not detectable | Not detectable | Not detectable |
| QES-61-020-W0-PE | 7 | Not detectable | Not detectable | Not detectable | Not detectable |
| QES-61-026-W0-PE | 2 | Not detectable | Not detectable | Not detectable | Not detectable |
| SAR-29-030 | 7 | Not detectable | Not detectable | Not detectable | 1.25E+00 |
| SAR-29-048 | 10 | Not detectable | Not detectable | Not detectable | 2.10E+00 |
| SAR-29-057 | 5 | Not detectable | Not detectable | Not detectable | Not detectable |

**Supp. Fig. 1. Quantification of human plasma samples after additional pelleting steps.** A. Schematic showing pelleting steps carried out on human plasma samples prior to quantification of bacterial copy number. B. Table showing bacterial copies detected in 11 human plasma samples subjected to additional pelleting. Where detectable, bacterial DNA was measured in fraction 4, corresponding to a high speed pellet. This demonstrates the bacterial DNA is not present in residual human cells (fraction 2) nor as free DNA in the supernatant (fraction 3).

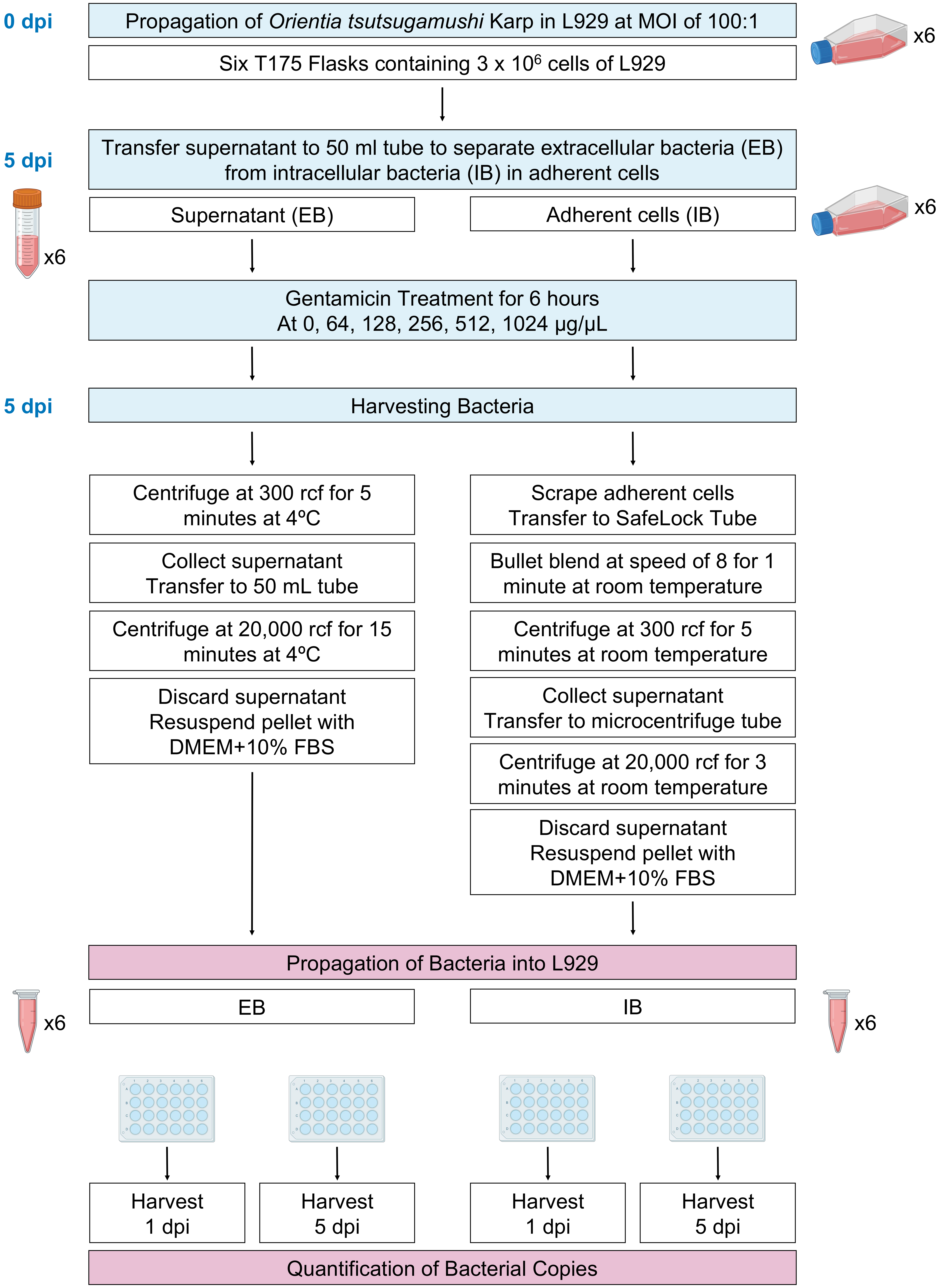


**Supp. Fig. 2. Overview of experimental workflow to quantify the effect of gentamicin treatment on IB and EB form Ot.**
